# Complex third-party effects in the *Dictyostelium*-*Paraburkholderia* symbiosis: prey bacteria that are eaten, carried, or left behind

**DOI:** 10.1101/2022.11.06.513053

**Authors:** Trey J. Scott, David C. Queller, Joan E. Strassmann

## Abstract

Symbiotic interactions may change depending on the abundance of third parties like predators, prey, or pathogens. Third-party interactions with prey bacteria are central to the symbiosis between *Dictyostelium discoideum* social amoeba hosts and *Paraburkholderia* facultative bacterial symbionts. Symbiosis with inedible *Paraburkholderia* allows host *D. discoideum* to carry prey bacteria through the dispersal stage where host amoebae aggregate and develop into fruiting bodies that disperse spores. Carrying prey bacteria benefits hosts when prey bacteria are scarce but harms hosts when prey bacteria are plentiful. Symbiont-carrying hosts leave some bacteria behind; this could explain the harm to hosts if left-behind bacteria include uneaten prey bacteria. Thus, understanding both benefits and costs in this symbiosis requires measuring how many prey bacteria are eaten, carried, and left behind by infected hosts. We found that *Paraburkholderia* infection makes hosts leave behind both symbionts and prey bacteria. However, the number of prey bacteria left uneaten was small and did not explain why infected hosts produced fewer spores than uninfected hosts. Turning to the bacteria that are carried, we found that hosts carry more prey bacteria after developing in prey- poor environments than in prey-rich ones. This suggests that carriage is actively modified to ensure hosts have prey in the harshest conditions. Our results show that multifaceted interactions with third parties shape the evolution of symbioses in complex ways.

## Introduction

The fitness effects of symbiotic interactions can change depending on the environment (Bronstein, 1994; Chamberlain *et al*., 2014; Horas *et al*., 2022; Scott *et al*., 2022). One crucial component of the environment that often selects for symbiotic interactions is a third species that interacts with hosts or symbionts (Palmer *et al*., 2008; Wendling *et al*., 2017; Wood *et al*., 2018; Hafer-Hahmann & Vorburger, 2020; Cassidy *et al*., 2022). For example, in the symbiosis between ants and *Acacia* plants, ants benefit *Acacia* by fending off herbivores. However, when herbivores were prevented from accessing *Acacias*, ordinarily mutualistic interactions between ants and *Acacias* shifted towards antagonism because ants no longer provided a benefit to host *Acacias* (Palmer *et al*., 2008). Such shifts can influence whether host and symbiont fitness interests are aligned or in conflict (Keeling & McCutcheon, 2017; Iwai *et al*., 2019). Fitness alignment or conflict then shapes the evolution of these interactions (Scott & Queller, 2019; O’Brien *et al*., 2021). However, the details of how third parties affect the fitness interests of symbiotic partners are not well understood for many kinds of symbioses (Chamberlain *et al*., 2014).

An important model of microbial symbiosis is the social amoeba *Dictyostelium discoideum*, that is famous for its complex and well-studied multicellular life cycle (Kessin, 2001; Jahan *et al*., 2021). *D. discoideum* is a soil amoeba that feeds on many bacteria (Raper, 1937; Brock *et al*., 2018). Upon starvation, amoebae aggregate to form a multicellular fruiting body that disperses spores to new soil patches (Kessin, 2001).

Development of the fruiting body progresses through well-studied stages starting with aggregation of individual amoebae which then become motile slugs (Medina *et al*., 2019). Slugs further develop into a fruiting body. During fruiting body development, about 20% of the cells die and become stalk while the remaining cells develop into spores that sit atop the stalk in a structure called a sorus (Strassmann & Queller, 2011).

*D. discoideum* has symbioses with certain facultatively intracellular *Paraburkholderia* bacteria (DiSalvo *et al*., 2015; Brock *et al*., 2020). In natural isolates, around 25% of wild-collected *D. discoideum* are infected (Haselkorn *et al*., 2019). There are three symbiont species – *P. agricolaris*, *P. hayleyella*, and *P. bonniae* – and we will refer to them collectively in this paper as either symbionts or simply *Paraburkholderia* (though readers should understand that there are many species of *Paraburkholderia* not involved in this symbiosis).

These symbionts are adapted for interacting with amoebae (Shu *et al*., 2018a; b; Brock *et al*., 2020). Symbionts move towards hosts (Shu *et al*., 2018b) and make host phagosomes less acidic so that the symbionts are not digested (Tian *et al*., 2022). *P. hayleyella* and *P. bonniea* also have reduced genomes relative to *P. agricolaris* and most other *Paraburkholderia* (Brock *et al*., 2020; Noh *et al*., 2022) and such reduction is often found in endosymbionts and pathogens that have been associated closely with their hosts during a long evolutionary history (McCutcheon & Moran, 2012). Some *D. discoideum* hosts may also be adapted to their symbionts. Hosts infected by *P. hayleyella* appear to suffer less from *P. hayleyella* toxicity than hosts that are naturally infected by *P. agricolaris* (Shu *et al*., 2018a).

Fitness of *D. discoideum* hosts and *Paraburkholderia* symbionts is affected by interactions with a third set of organisms, the various prey bacteria that are eaten by host amoebae (Brock *et al*., 2011, 2018; DiSalvo *et al*., 2015; Scott *et al*., 2022). *Paraburkholderia* symbionts are largely inedible by hosts but they cause host carriage of additional, edible bacteria (Brock *et al*., 2011; DiSalvo *et al*., 2015; Khojandi *et al*., 2019). Edible prey bacteria are mostly carried extracellularly in the sorus while *Paraburkholderia* symbionts are mostly carried inside host spores (Khojandi *et al*., 2019). When host spores disperse to locations where prey bacteria are rare, they benefit from *Paraburkholderia* infection because they are able to seed new prey bacteria populations (Brock *et al*., 2011; DiSalvo *et al*., 2015; Scott *et al*., 2022). When hosts disperse to prey-rich conditions, infected hosts do not gain from carrying prey bacteria and instead experience a cost producing fewer spores relative to uninfected *D. discoideum* (Brock *et al*., 2011; DiSalvo *et al*., 2015). Higher *Paraburkholderia* densities impose higher costs on hosts but again this is affected by third-party prey, with the effect being stronger in pre-rich conditions (Scott *et al*., 2022).

The third-party bacteria can also affect *Paraburkholderia* fitness due to competition (Medina *et al*., 2023). One consequence is that symbionts reach higher densities when they are carried to locations with fewer prey bacteria (Scott *et al*., 2022). Thus prey-poor conditions for the host are low-competition conditions for the symbiont (though for consistency, we will stick to calling them prey-poor and prey-rich conditions rather than low-competition and high-competition conditions).

While the importance of prey bacteria has been well established for this symbiosis (DiSalvo *et al*., 2015; Khojandi *et al*., 2019; Scott *et al*., 2022), there are still many questions about the effects of prey bacteria during development of *D. discoideum*. In this study, it is the disposition of prey bacteria during host development that is of interest – should they be eaten, carried, or left behind?

Leaving prey bacteria behind would not seem to be adaptive, but this appears to happen when hosts are infected by *Paraburkholderia* (Brock *et al*., 2011, 2016b).

Initially, before the distinct roles of inedible symbionts and edible prey were appreciated, bacteria left behind on plates was taken as a possible sign of “prudent predation” by host amoebae — if some bacteria were going to be saved for carriage, hosts cannot eat all the available prey and may stop feeding and start developing earlier (Brock *et al*., 2011). It was hypothesized that hosts leaving prey bacteria uneaten might explain why carriage is costly in some conditions (Brock *et al*., 2011). Hosts that were prudent predators left prey uneaten and missed out on potential growth and proliferation.

The essential role of *Paraburkholderia* in the symbiosis, including causing hosts to leave behind bacteria is now known (DiSalvo *et al*., 2015; Brock *et al*., 2016b). At least some of the bacteria left behind are not prey bacteria but inedible *Paraburkholderia* symbionts (Scott *et al*., 2022). Because of this new information, the prudent predation hypothesis should be re-evaluated by determining the relative abundances of left behind prey bacteria and *Paraburkholderia* symbionts. The fitness consequences of leaving prey behind, in terms of spore production and development time, should also be evaluated.

While infected hosts may leave some prey bacteria uneaten, they also gain the ability to carry prey bacteria along with the dispersing host spores in sori (Brock *et al*., 2011; DiSalvo *et al*., 2015; Khojandi *et al*., 2019). It is unknown whether the number of carried prey bacteria changes in different environments. If carriage is passive, we expect hosts to carry more prey after developing in a prey-rich environment and fewer prey bacteria after developing in a prey-poor environment.

Alternatively, the ability to carry prey bacteria could reflect the evolutionary interests of hosts and symbionts. Since soil environments tend to be spatially and temporally structured (Sun *et al*., 2003; Vos *et al*., 2013), developing in a prey-poor environment may be associated with an increased probability that the next environment will also tend to be prey-poor. If this is the case, it would be more adaptive for hosts if they carried more prey bacteria after developing in a prey-poor environment than in a prey-rich environment. This may also be adaptive for the *Paraburkholderia* symbionts since it allows better survival of their hosts.

We investigate three questions about the complex role of prey bacteria in the symbiosis between *D. discoideum* and two commonly studied *Paraburkholderia* symbionts, *P. agricolaris* and *P. hayleyella* (Shu *et al*., 2018a; Garcia *et al*., 2019; Scott *et al*., 2022; Medina *et al*., 2023). We first re-evaluate some of the ideas behind the prudent predation hypothesis. Second, we ask whether any prey bacteria that are left uneaten results in costs due to delayed development or reduced spore production. Lastly, we turn to whether the number of prey bacteria carried inside sori changes between prey-poor and prey-rich environments.

## Methods

### Clones and Culturing Methods

In our experiments, we used several types of infected and uninfected hosts (Table 1). We used two types of infected host, naturally infected hosts (n=7 for each symbiont species) and, to better control for infection levels, cured-and-reinfected hosts (n=4 for each symbiont species). The naturally infected hosts were compared to naturally uninfected controls (n=6) and the cured-and-reinfected host were compared to a different set of naturally uninfected controls (n=4). To better control for genetics, cured-and- reinfected hosts were also compared to their cured clonemates (n=4 for each symbiont species). To generate cured-and-reinfected clones, naturally infected clones were cleared of their natural infections with antibiotics and then reinfected at 1:1000 ratios of *Paraburkholderia* to prey bacteria to control infection densities (Scott *et al*., 2022). Data reported here for cured-and-reinfected hosts and their uninfected controls come from experiments performed in Scott et al. (2022) but here we include additional measures not included in the original study. Naturally uninfected controls in this comparison were also treated with antibiotics to control for antibiotic treatment. We stored all prepared clones in a -80 C freezer for repeated use.

**Table 1:** Clones used in experiments.

| Infection Type | Description | *Paraburkholderia*<br>Treatment | Clones |
| --- | --- | --- | --- |
| Cured-and-reinfected | Clones that were cured of any native symbionts with tetracycline and reinfected in the lab with 0.1% of their native symbiont | *P. agricolaris*<br>infected | QS159,<br>QS161,<br>QS606, NC21 |
|  |  | *P. hayleyella*<br>infected | QS395, QS45,<br>QS38, QS23 |
| Naturally uninfected controls<br>(for comparison with cured-and-reinfected) | Control clones that were treated with tetracycline. | Naturally uninfected control | QS6, QS138,<br>QS472, QS527 |
| Cured (for same-clone comparison) | Clones that were cured of any native symbionts with | Cured of *P. agricolaris*<br>infection | QS159,<br>QS161,<br>QS606, NC21 |
| with cured-and-reinfected) | tetracycline, but not reinfected. | Cured of *P. hayleyella* infection | QS395, QS45, QS38, QS23 |
| Naturally infected hosts | Clones with natural infections that have not been treated with antibiotics. | *P. agricolaris* infected | QS494, QS756, QS788, QS113, QS453, QS70, QS606 |
|  |  | *P. hayleyella* infected | QS45, QS101, QS46, QS2, QS23, QS529, QS38 |
| Naturally uninfected controls (for comparison with naturally infected) | Clones without natural infections that have not been treated with antibiotics. | Naturally uninfected control | QS4, QS6, QS14, QS18, QS9, QS8 |

To remove any effects of being in the freezer and to ensure that infected amoebae carried *K. pneumoniae* prey bacteria, we grew amoebae through one round of feeding and fruiting body formation and then collected spores to initiate our experiments. This step may allow infection densities time to equilibrate (Miller *et al*., 2020). Amoebae were grown from frozen spores at room temperature on 100 x 15mm plates filled with 30mL SM/5 agar (2 g glucose (Fisher Scientific), 2 g Bacto Peptone (Oxoid), 2 g yeast extract (Oxoid), 0.2 g MgSO_4_ * 7H_2_O (Fisher Scientific), 1.9 g KH_2_PO_4_ (Sigma-Aldrich), 1 g K_2_HPO_4_ (Fisher Scientific), and 15 g agar (Fisher Scientific) per liter) with 200 μL of 1.5 OD_600_ fluorescently labeled *K. pneumoniae* suspended in KK2 buffer (2.25 g KH_2_PO_4_ (Sigma-Aldrich) and 0.67 g K_2_HPO_4_ (Fisher Scientific) per liter) spread over the entirety of the agar. *K. pneumoniae* used in this study expressed green-fluorescent protein (GFP) and were provided by dictyBase (Fey et al. 2019).

### Dispersal experiment setup

To determine the consequences of hosts leaving behind bacteria, we performed a dispersal experiment to seed new plates with sorus contents. To do this, we transferred 200 μL of 2x10^5^ spores/mL suspension obtained from the equilibration step to plates with or without 200 μL of 1.5 OD_600_ GFP-expressing *K. pneumonia*e. Spore counts to make spore suspensions were obtained using a hemocytometer. OD_600_ for plating bacteria was measured with a BioPhotometer (Eppendorf, New York). We let bacteria and amoebae on plates proliferate at room temperature for six days, enough time for amoebae to form fruiting bodies.

We replicated these dispersal experiments with cured-and-reinfected hosts along with uninfected controls on two separate dates. To understand if our results held with natural infections, we performed an additional dispersal experiment with naturally infected clones and uninfected controls. To rule out that leaving uneaten prey bacteria was a host trait, we also performed a separate set of dispersal experiments comparing cured clones to their reinfected counterparts.

### Measurement of bacteria and host spore production

After dispersed amoebae and bacteria were allowed to proliferate for six days (to allow hosts to complete development into fruiting bodies), we measured the density of uneaten prey bacteria, *Paraburkholderia*, and host spore production. To measure the density of uneaten prey bacteria, we first collected host spores and bacteria from after the dispersal step. We collected host spores and bacteria by washing plates with 15 mL of KK2 buffer after six days of growth. We counted host spores from this washed solution using a hemocytometer. To measure bacterial density, we first removed host spores by centrifuging the wash solution for three minutes at 360 rcf and reserving the supernatant. We measured the optical density of the supernatant at 600 nm (OD_600_) as well as fluorescence with an excitation wavelength of 485 and emission wavelength of 515 nm in a 96 well plate with a Tecan Infinite 200 Pro microplate reader. Removing host spores by manually removing sori with a pipette tip resulted in similar densities of uneaten prey bacteria consistent earlier findings (Khojandi *et al*., 2019) that the number of prey bacteria inside spores and sori is minimal relative to that left on the plate (data not shown).

Since the supernatant from infected clones contains GFP-expressing *K. pneumoniae* and unlabeled *Paraburkholderia*, the total OD_600_ is due to both kinds of bacteria but fluorescence comes only from *K. pneumoniae*. To calculate the amount of OD_600_ due to fluorescing *K. pneumoniae*, we generated a standard curve relating fluorescence to OD_600_ using serial dilutions of GFP-expressing *K. pneumoniae* in KK2 and used this curve to predict the OD_600_ of *K. pneumoniae* in our samples. The remaining OD_600_ is the amount due to *Paraburkholderia* symbionts, after also subtracting off the background OD_600_ of the KK2 buffer. An OD_600_ of 0.1 translates to around 5 x 10^7^ *K. pneumoniae* cells or 1 x 10^8^ *Paraburkholderia* cells according to the validation dataset in Scott et al. (2022).

### Development assays for cured and reinfected hosts

To determine how symbionts and their interactions with prey affected host development time, we took time-lapse images of cured hosts and reinfected counterparts developing in six-well plates filled with 10 mL SM/5. Each 6-well plate contained three wells with cured hosts and three wells with reinfected hosts of either *P. hayleyella* clones (QS23, QS38, and QS45) or *P. agricolaris* clones (NC21, QS159, and QS161). We repeated development experiments on six-well plates three times for each *Paraburkholderia* species. We first grew clones from the freezer on SM/5 plates for six days in case there were freezer effects, as above. We then collected host spores and plated 30 μL of 2x10^5^ spores per mL and 30 μL of 1.5 OD_600_ *K. pneumoniae* in each well. Photos were taken every hour until fruiting bodies developed using a Canon EOS Mark IV. We inspected photos to determine time points for when aggregates, slugs, and fruiting bodies first appeared in each well.

### Measurement of carried bacteria

To determine how prey context affected the number of prey bacteria carried in sori, we used cured-and-reinfected hosts that were grown on prey-poor and prey-rich plates after the dispersal step. We haphazardly sampled a single sorus from each of our experimental plates from reinfection experiments (replicated on two different dates). To count the number of carried prey, we suspended single-sorus contents in KK2 buffer and plated out serial dilutions from no dilution to 1:1000 dilutions. Since *K. pneumoniae* were labeled with GFP, we could differentiate colonies of *K. pneumoniae* and *Paraburkholderi*a. We counted colonies from the dilution plate that appeared closest to having 100 colonies and then back-calculated to get the total count for a single sorus.

This experiment was completely replicated on two separate dates.

### Statistical Methods

To understand how *Paraburkholderia* infection affected the density of *K. pneumoniae* bacteria left uneaten, we fit linear models in R (version 3.6.3). Because experiments with cured-and-reinfected (and uninfected control) hosts were performed on two different dates, we used linear mixed models (LMM) with date as a random effect. We fit LMMs using the nlme4 package (Pinheiro & Bates, 2006) in R. For other comparisons that involved only single measures, we used linear models fit by ordinary linear regression (LM) with the lm function in R. We performed Tukey post-hoc tests for pairwise comparisons using the *emmeans* package (Lenth *et al*., 2019).

To determine whether the timing of development was affected by *Paraburkholderia* infection, we used generalized linear mixed models (GLMMs) with Poisson errors. We included the date that experiments were performed as a random effect to capture variation within plates.

To test how prudent predation affects host spore production relative to other explanations, we again used LMMs and LMs for cured-and-reinfected and naturally infected infections, respectively, along with respective uninfected controls. If prudent predation reduces host spore production, we expected a decrease in spore production with increasing uneaten prey bacteria. Differences in host fitness could also be affected by the density of *Paraburkholderia* left on plates (Scott *et al*., 2022) or by infection category (uninfected vs infected) (Brock *et al*., 2011; DiSalvo *et al*., 2015; Haselkorn *et al*., 2019). To account for these possibilities, we also fit models with these variables along with a model that includes both uneaten prey density and *Paraburkholderia* density. In total, we compared five models of spore production: (1) uneaten prey density (Prey model), (2) *Paraburkholderia* density (Para model), (3) uneaten prey density + *Paraburkholderia* density (Prey + Para model), (4) categorical infection status (Infection model), and (5) a null model fit with only the intercept for each *Paraburkholderia* species.

To compare effects more easily across the different models, we scaled all variables by subtracting the mean and dividing by twice the standard deviation. This scaling procedure is useful for comparing models with multiple continuous predictors (density of prey bacteria and symbionts) and with categorical factors (infected and uninfected) because all measures end up on roughly the same scale (Gelman *et al*., 2020).

To compare the fit of the five models described above, we calculated AICc for each model. AICc is a measure of model fit that accounts for model complexity and that corrects for small sample sizes (Burnham & Anderson, 2004). Models that fit the data best have lower AICc values. For our model selection analysis we show results with 85% confidence intervals instead of the standard 95% confidence intervals because 85% confidence intervals are more consistent with model selection with AIC (Arnold, 2010). To measure potential collinearity in our models that included prey and *Paraburkholderia* density, we calculated variance inflation factors (VIF) using the check_collinearity function in the performance package (Lüdecke *et al*., 2021). VIFs lower than 5 indicate low collinearity, values between 5 and 10 indicate moderate collinearity, and values above 10 indicate high collinearity.

To compare the ability of hosts to carry *K. pneumoniae* inside fruiting bodies in prey-rich and prey-poor environments, we used a zero-inflated negative binomial model (ZINB) fit with the glmTMB package (Brooks *et al*., 2017) in R. This zero-inflated model incorporated two processes: (1) zero-inflation for whether fruiting bodies contained or did not contain prey bacteria and (2) negative binomial for the counts of carried prey bacteria. The zero-inflation parameter captures whether hosts are more likely to carry while the negative binomial parameter captures whether hosts carry more prey bacteria when they carry. Using this ZINB, we compared different hypotheses about how prey-rich and prey-poor environments affected the ability to carry and the number carried with AICc. We fit models to carriage data from both species and asked whether models including prey environment, *Paraburkholderia* species, or the interaction between prey environment and species had low AICc values. We did this model selection for both the zero-inflation part of the model and for the negative binomial part of the model. We also compared models with the random effects of clone and date included in either the zero-inflation or negative binomial parts of the model.

## Results

### D. discoideum hosts infected with Paraburkholderia leave some prey bacteria uneaten

We investigated the prey bacteria left behind after hosts formed fruiting bodies by measuring the density of leftover *K. pneumoniae* from cured-and-reinfected hosts compared to naturally uninfected controls. These prey densities are estimated from fluorescence measurements since the *K. pneumoniae* used in this study expresses green fluorescent protein (GFP; see methods). First, we wanted to confirm that some of the bacteria hosts leave behind are prey bacteria and not just *Paraburkholderia* symbionts. Leaving prey bacteria uneaten so that hosts miss out on potential growth and proliferation was a key component of the prudent predation hypothesis, but previous studies did not differentiate what kind of bacteria were left behind (Brock et al. 2011, 2016*b*).

Cured-and-reinfected hosts with *P. agricolaris* left 4.4 times as much prey bacteria as naturally uninfected hosts (LMM; estimate = 0.009, se = 0.003, df = 20, P < 0.005; Figure 1A). Cured-and-reinfected hosts that were infected with *P. hayleyella* left 7.4 times as much prey bacteria than naturally uninfected hosts (Figure 1A; LMM; estimate = 0.017, se = 0.003, df = 20, P < 0.001).

**Figure 1:**
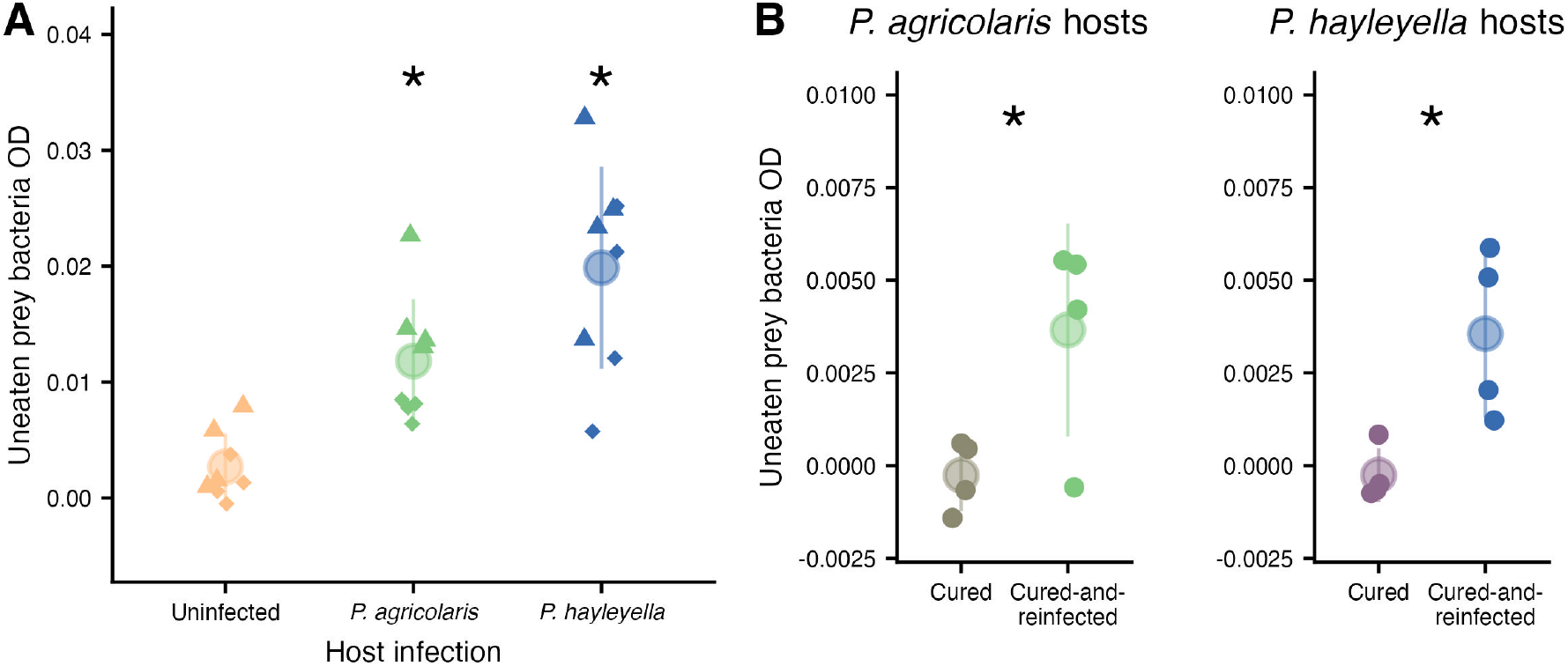
Symbionts cause hosts to leave prey uneaten. (A) Density left on plate of uneaten *K. pneumoniae* prey bacteria (measured by OD_600_) for naturally uninfected hosts, *P. agricolaris* hosts that have been cured and then reinfected, and *P. hayleyella* hosts that have been cured and then reinfected. (B) Density left on plate of uneaten prey bacteria for cured *P. agricolaris* and *P. hayleyella* hosts compared to their cured-and-reinfected counterparts. Dot and lines show mean and standard deviation, respectively. Asterisks indicate significant differences (in panel A comparisons are between infected and uninfected). Point shapes of small points indicate different dates on which experiments were replicated.

This comparison supports the view that symbionts cause prey bacteria to be left uneaten. To control for host clone genotypes, we next compared cured-and-reinfected hosts to the same host genotypes that were cured of their native *Paraburkholderia* symbionts but not reinfected. Cured-and-reinfected hosts left more prey bacteria than their cured counterparts (*P. agricolaris*: LM; estimate = 0.004, se = 0.002, df = 6, P = 0.041, *P. hayleyella*: LM; estimate = 0.004, se = 0.001, df = 6, P = 0.019; Figure 1B).

### Host development time is not affected by Paraburkholderia infection

We hypothesized that leaving prey bacteria uneaten would result in faster starvation and faster development by infected hosts. To test this, we took time-lapse photos of amoebae aggregating into fruiting bodies, comparing cured-and-reinfected hosts to the same host genotypes that were cured of their *Paraburkholderia* symbionts.

We determined time courses for when hosts aggregated, formed slugs, and formed fruiting bodies. We were unable to detect that development times were slower for cured- and-reinfected hosts relative to cured hosts for *P. agricolaris* (GLMM; estimate across all developmental timepoints = 0.015, se = 0.037, P = 0.682) or *P. hayleyella* (GLMM; estimate across all developmental timepoints = 0.044, se = 0.037, P = 0.231; Figure 2A&B). Individual developmental stages also did not differ due to *Paraburkholderia* infection (P > 0.05).

**Figure 2:**
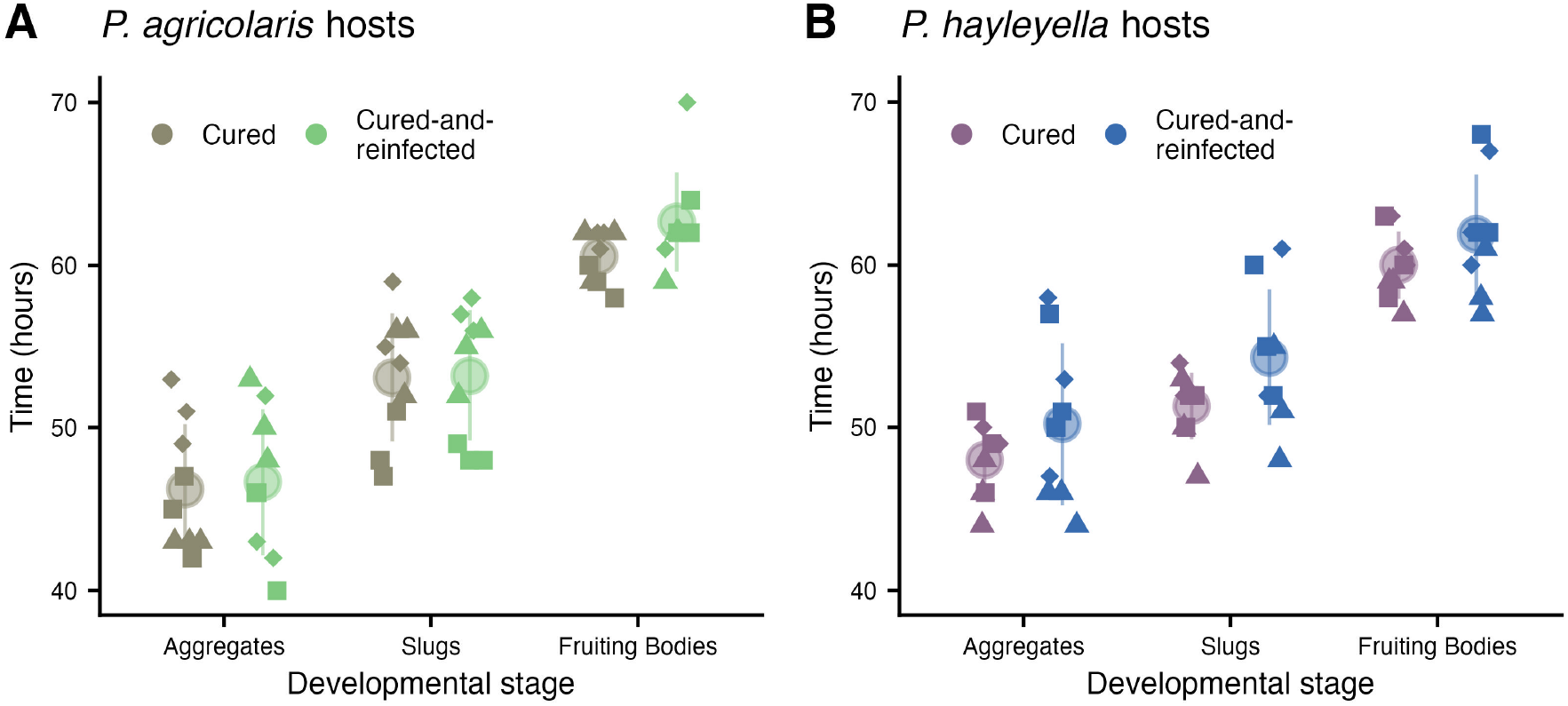
Symbionts do not affect developmental time. Developmental time points at different stages of development for cured hosts and their counterparts that have been cured and then reinfected with (A) *P. agricolaris* and (B) *P. hayleyella* hosts. Dot and lines show mean and standard deviation, respectively. Point shapes of small points indicate different dates on which experiments were replicated.

### Prey bacteria left on plates are not associated with reduced spore production in hosts

Leaving more prey uneaten is suspected to lower host fitness and could therefore explain the cost of *Paraburkholderia* infection relative to uninfected hosts (Brock *et al*., 2011). Alternately, the cost of infection may be more directly due to *Paraburkholderia*, either infection itself or the density of *Paraburkholderia* symbionts. In addition to these first three models (food bacteria density, *Paraburkholderia* infection status, and *Paraburkholderia* density), we also fit a model that included both uneaten prey bacteria density and *Paraburkholderia* density. To compare the relative support for these different hypotheses, we fit a model for each hypothesis (see Methods) and compared fitted models using AICc.

We first tested the role of uneaten prey bacteria and *Paraburkholderia* left on plates on host spore production using our cured-and-reinfected hosts and naturally uninfected controls (Table 1). For infections with either species of *Paraburkholderia*, null models fit to only the intercept fit the data best (Figure 3A & S1; the model that includes uneaten prey bacteria and *Paraburkholderia* density for *P. agricolaris* cured-and-reinfected hosts — Prey + Para — did show confidence intervals that did not overlap zero, but this was the worst model in terms of AICc). We thus found no evidence that leaving prey explained the cost of infection for cured-and-reinfected hosts relative to uninfected controls.

**Figure 3:**
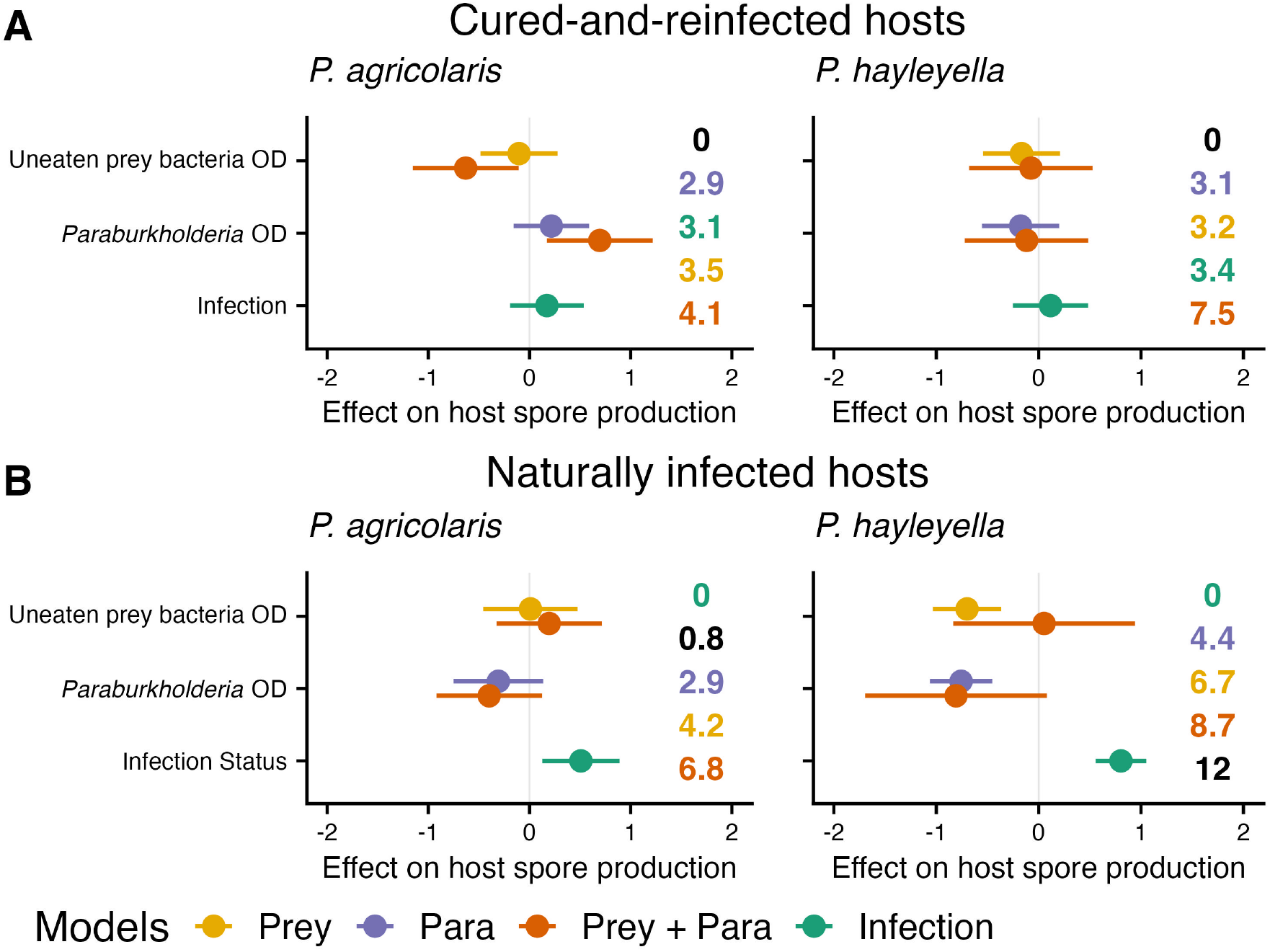
Host spore production as a function of different bacteria densities on plates and infection status in (A) cured-and-reinfected (B) and naturally infected hosts relative to their respective uninfected counterparts (see Table 1). We compared models of host spore production (shown in different colors), predicted by *Paraburkholderia* density only, uneaten prey bacteria only, both *Paraburkholderia* and uneaten prey bacteria, or a categorical variable for infection status (infected with *Paraburkholderia* or not). Estimated effects are shown as points and 85% confidence intervals are shown as lines (null models with only the intercept are not shown). ΔAICc values are shown in increasing order from top to bottom and colored according to the kind of model. Intercepts are not shown.

Naturally infected hosts and their uninfected controls also showed no support for a cost due to leaving prey bacteria behind. Models of host spore production that included uneaten prey (Prey model in Figure 3B) were poor predictors of host spore production.

This poor performance of prey as a predictor was also true when *Paraburkholderia* was included as a covariate (Prey + Para model). This lack of fit for the *P. hayleyella* Para + Prey model may be due to correlations between *Paraburkholderia* density and uneaten food bacteria (VIF = 7.62).

These results show no support for a cost due to leaving prey bacteria behind (even when accounting for *Paraburkholderia* density). We next turned to the direct role of *Paraburkholderia*. Because models fit to only the intercept were the best for cured-and- reinfected hosts relative to their controls, there was no evidence that *Paraburkholderia* was costly in these clones, the same result as previously shown (Scott *et al*., 2022). In contrast, naturally infected hosts were afflicted by a cost of *Paraburkholderia* infection (Figure 3B & S2). The best models for both *P. agricolaris* and *P. hayleyella* showed that infected hosts had lower spore production than uninfected hosts (*P. agricolaris*: LM; estimate = -0.508, se = 0.247, df = 11; *P. hayleyella*: LM; estimate = -0.804, se = 0.160, df = 11). *Paraburkholderia* density predicted host spore production less well than infection status. However, models that included *Paraburkholderia* density on its own were better than models with prey bacteria density (Figure 3B & S2). These models thus show little evidence for a cost of uneaten food bacteria and instead show that *Paraburkholderia* infection is responsible for reduced host spore production in naturally infected hosts.

### Uneaten prey bacteria are a minority of left-behind bacteria

Prey bacteria left behind did not have the predicted effects on either development time or spore production costs. One reason may be that the amount of left behind prey is not as great as formerly thought (Brock *et al*., 2011). Hosts may leave too few prey bacteria to noticeably affect spore production. In fact, uneaten prey bacteria make up a minority of left-behind bacteria. Uneaten prey bacteria are about 9% and 14% of bacteria on plates for *P. agricolaris* and *P. hayleyella* cured-and-reinfected hosts, respectively, and 21% and 19% for *P. agricolaris* and *P. hayleyella* naturally infected hosts, respectively (Figure 4).

**Figure 4:**
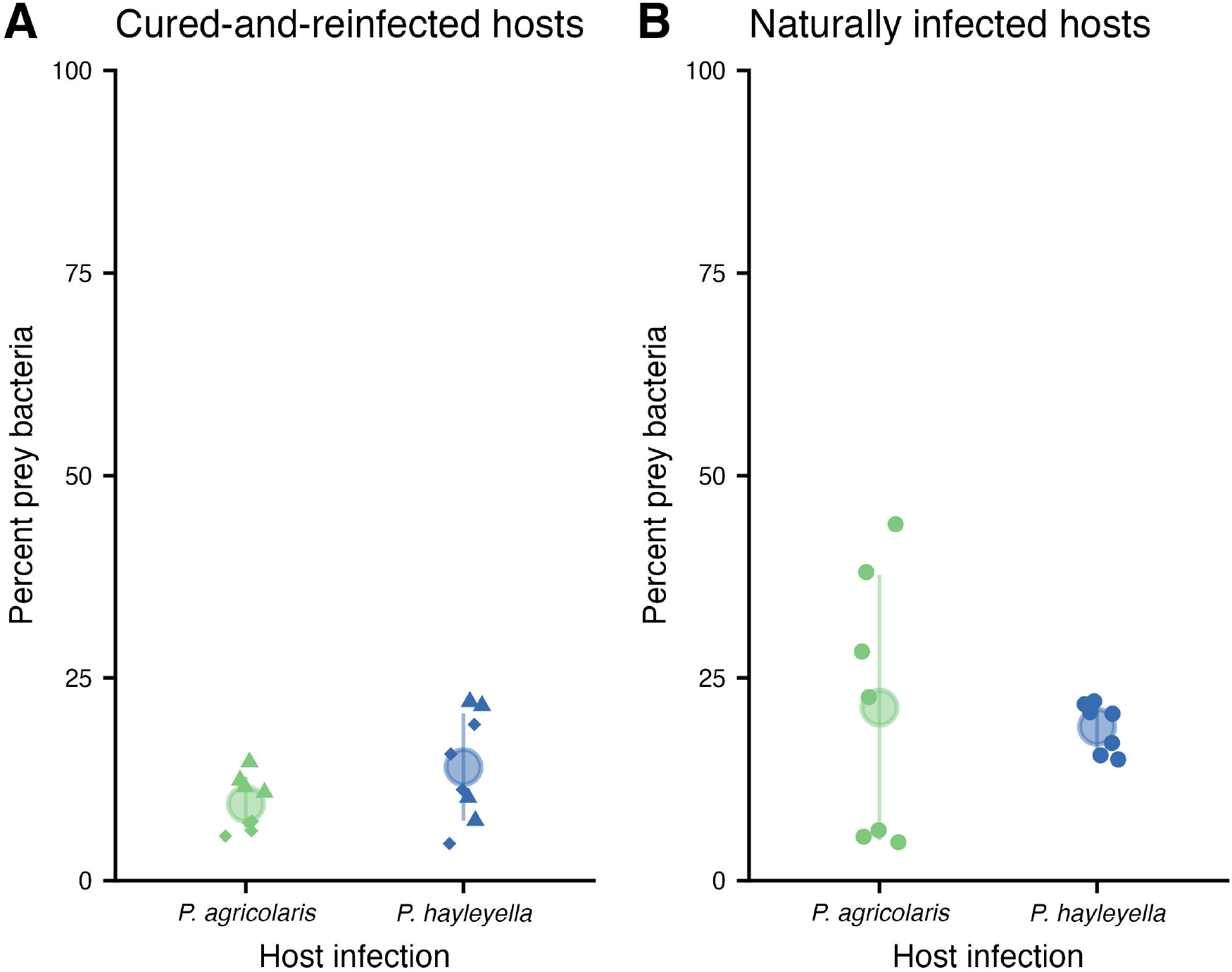
Most of the bacteria left behind by infected hosts were not prey bacteria. Percent of the total left-behind bacteria (includes prey bacteria and *Paraburkholderia* symbionts) that was prey bacteria from cured-and-reinfected (A) and naturally infected (B) hosts. Dot and lines show mean and standard deviation, respectively. Point shapes for small points indicate different dates on which experiments were replicated

### More prey bacteria are carried after hosts develop in prey poor conditions

To determine whether the number of prey bacteria carried in sori simply reflects the number of prey bacteria in the previous environment or is modified according to the interests of hosts or symbionts, we measured the number of fluorescent prey bacteria inside sori after growth of cured-and-reinfected hosts on prey-rich and prey-poor plates (Figure 5A). We found that for both *Paraburkholderia* species (Figure 5B&C), sori produced in prey-rich environments were less likely to contain prey bacteria than those in prey-poor environments (ZINB, zero-inflation estimate = 1.950, se = 0.987). This shows that the ability to carry prey bacteria is affected by prey context, but in the opposite direction to that expected if prey was carried according to the density of prey bacteria in the environment. We did not detect a difference in the number of prey bacteria that were caried between prey-rich and prey-poor environments (Figure 5D&E). Thus, the ability to carry prey bacteria may depend on the fitness interests of hosts and symbionts that benefit from carrying prey bacteria when harsh conditions are expected.

**Figure 5:**
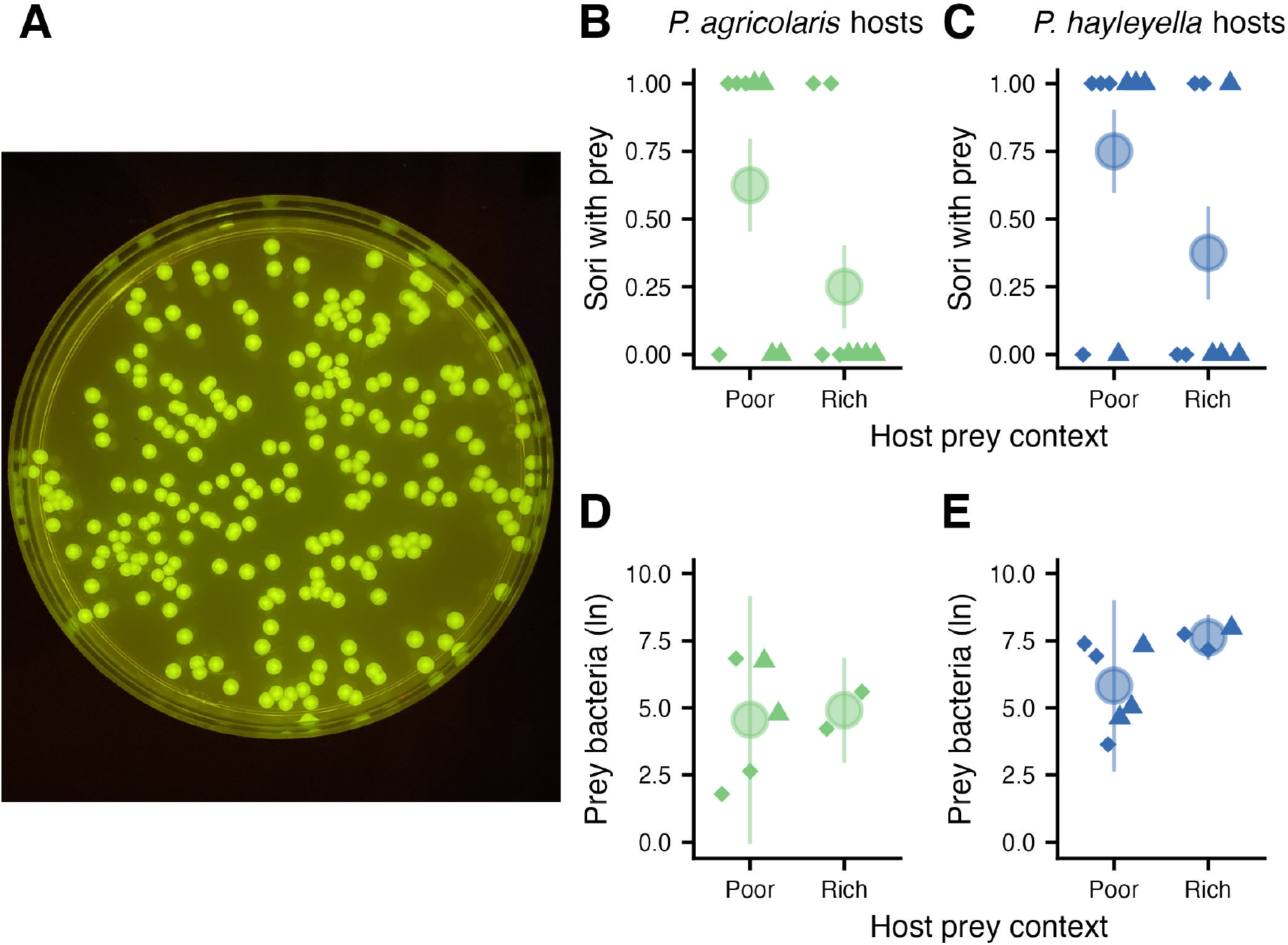
Hosts are less likely to carry prey bacteria in sori after developing in prey-rich environments. (A) Photo of fluorescent *K. pneumoniae* prey bacteria colonies plated from an individual sorus. (B&C) Sori with carried food bacteria for cured-and-reinfected *P. agricolaris* (B) or *P. hayleyella* (C) hosts from prey-poor and prey-rich contexts. Carriage outcomes for individual fruiting bodies are shown as small points that indicate carried prey (1) or did not carry prey (0). Large points and lines show the proportion of sori with prey and the standard deviation. (D&E) Number of prey bacteria of sori that contained prey (1s in B&C), for hosts carrying *P. agricolaris* (D) or *P. hayleyella* (E) from prey-poor and prey-rich contexts. Large points and lines in D&E show the average and standard deviation. Carried prey bacteria were quantified by plating serial dilutions (from undiluted to 1:1000) of single fruiting bodies. Point shapes for individual fruiting bodies (small points) indicate different dates on which experiments were replicated.

## Discussion

Third parties that interact with symbiotic partners commonly affect the outcome of symbiotic interactions (Palmer *et al*., 2008; Chamberlain *et al*., 2014; Hafer-Hahmann & Vorburger, 2020; Cassidy *et al*., 2022) and alter selection for cooperation and conflict between symbiotic partners (Keeling & McCutcheon, 2017; Wood *et al*., 2018). Often the details of how third parties affect fitness effects are unknown. In the symbiosis between *D. discoideum* social amoebae and *Paraburkholderia* bacteria, the third-party prey bacteria are normally eaten by the host but can also be carried or left behind. We studied how many prey bacteria are carried and left behind, and their impacts, by tracking fluorescently labeled prey bacteria (*K. pneumoniae*) during fruiting body formation by hosts.

Our most surprising finding was that infected hosts were more likely to carry prey bacteria after growing in prey-poor environments (Figure 5). This result is surprising because it means that hosts or symbionts actively change the number of carried prey bacteria depending on environmental conditions. This result provides possible evidence for mutual benefit between *D. discoideum* hosts and *Paraburkholderia* symbionts. Hosts and symbionts are likely to benefit from modifying carriage in this way if future soil environments tend to resemble past soil environments. Repeated prey-poor environments are where hosts should most benefit from carrying and seeding out prey bacteria.

*Paraburkholderia* symbionts could also benefit from carriage of prey because keeping hosts alive means there are hosts to further disperse *Paraburkholderia*. However, this dispersal benefit to the symbionts must be balanced against the costs of increased competition with the prey bacteria.

We speculate that carrying more prey in prey-poor contexts may represent cooperation between hosts and symbionts that allows the symbiosis to persist over repeated harsh environments. Such harsh conditions are potentially an important force shaping cooperation in this (Scott *et al*., 2022, 2023b) and other symbioses (Henry *et al*., 2021; Veresoglou *et al*., 2021; Cornwallis *et al*., 2023). These studies in symbiotic systems complement research on the role of harsh environments in the evolution of cooperation that has been focused on interactions between members of the same species (Kennedy *et al*., 2018), most commonly in cooperatively breeding birds (Cornwallis *et al*., 2017; Griesser *et al*., 2017; Capilla-Lasheras *et al*., 2021).

Although seeding out prey bacteria has been shown to be important when we move spores to a prey-poor environment in the lab, we have little idea how frequently this occurs in nature. The finding that carriage of prey bacteria changes in an apparently adaptive fashion suggests that it is indeed important in the field. The importance of third-parties in this case may be an interesting exception to the relatively small effects from third-parties that have been observed across other studies (Chamberlain *et al*., 2014).

Questions remain about how hosts and symbionts affect the carriage of prey bacteria. *Paraburkholderia* are more often carried inside spores while prey bacteria are more often carried outside spores (Khojandi *et al*., 2019). Differences in carriage of bacteria could potentially result from differences in lectins (Dinh *et al*., 2018) or polyphosphates (Rijal *et al*., 2020) that affect the ability of bacteria to invade and survive inside hosts. Hosts in the absence of their *Paraburkholderia* symbionts are able to modify the contents of fruiting bodies through their immune cells that protect against toxins and bacteria by collecting potential threats and dropping off during the slug stage (Chen *et al*., 2007). However, the role of these immune cells in the symbiosis (Brock *et al*., 2016a; Scott *et al*., 2023a) needs to be further explored. Manipulation of phagosomes could also play a role in determining the contents of fruiting bodies. *Paraburkholderia* symbionts increase the pH of phagosomes, presumably to prevent host digestion of symbionts (Tian *et al*., 2022). Similar modification of lysosomes is used by pathogens to evade human immune cells during infection (Isberg *et al*., 2009; Leseigneur *et al*., 2020). More work is needed to understand how symbionts and food bacteria get into fruiting bodies and how both hosts and symbionts contribute to bacterial carriage.

We also found that *Paraburkholderia* infection prevents hosts from eating all the prey bacteria in an environment (Figure 1). A prior study suggested that leaving prey bacteria uneaten because of prudent predation may explain the cost of infection relative to uninfected hosts (Brock *et al*., 2011). Instead, we found that the quantity of left-behind prey bacteria may be too small to noticeably affect host spore production since we observed that only a minority of left-behind bacteria was food bacteria, with the majority being indigestible *Paraburkholderia* (Figure 4).

To get a better sense of how important left-behind prey bacteria could be for hosts, we can use the density vs bacteria count curve in Figure S1B of Scott et al. 2022 and the average left-behind prey bacteria in this study to estimate the number of individual left-behind prey bacteria. Using this estimation, *P. agricolaris* infected hosts left 7.060x10^6^ individual prey bacteria and *P. hayleyella* infected hosts left 1.206x10^7^ individual prey bacteria. Using Kessin’s (2001, p. 21) rough estimate that an amoebae needs to eat 1,000 bacteria to divide, we calculated that the number of uneaten prey bacteria is only enough to produce 0.002% and 0.004% more spores than what we recovered from plates (collected spores were around 3x10^8^) for *P. agricolaris* or *P. hayleyella* infected hosts, respectively. These percentages are a rough and conservative approximation as they assume that amoebae devote all consumed prey bacteria to proliferation. However, the calculations show that it is unlikely that uneaten prey bacteria can affect host fitness in a detectable manner.

Instead of hosts paying a cost in potential growth because they leave prey bacteria uneaten, we suspect that *Paraburkholderia* infection causes both observations: that hosts leave prey uneaten and that infected hosts pay a cost. Tentative support for this idea comes from our findings that *Paraburkholderia* and uneaten prey bacteria densities were correlated. This correlation could result from higher *Paraburkholderia* densities interfering with the ability of hosts to eat prey bacteria.

An interesting remaining question in the *D. discoideum*-*Paraburkholderia* symbiosis and in other symbioses is the mechanisms that result in reduced fitness when hosts are infected. One potential explanation is that *Paraburkholderia* symbionts directly feed on host cells or otherwise extract nutrients from hosts. Measures of *Paraburkholderia* density inside sori have so far not been found to be correlated with host fitness within species (Miller *et al*., 2020), though the more toxic *P. hayleyella* does appear to infect a higher percentage of spores than less toxic *P. agricolaris* (Shu *et al*., 2018a; Khojandi *et al*., 2019). Another promising hypothesis that deserves further study is that hosts have lower spore production because of “collateral damage” from competitive interactions between *Paraburkholderia* and prey bacteria (Scott *et al*., 2022). Competition between bacteria is often mediated by chemical warfare (Granato *et al*., 2019) that could reduce host *D. discoideum* fitness as a side-effect.

The role of *Paraburkholderia* in reducing host spore production is more evidence in support of there being some evolutionary conflict in this symbiosis (DiSalvo *et al*., 2015; Scott *et al*., 2022). However, the magnitude of conflict may differ across *Paraburkholderia* symbiont species. This conflict may be more pronounced between *P. hayleyella* and its hosts than between *P. agricolaris* and its hosts. We found that *P. hayleyella* density could explain some of the decrease in naturally infected host spore production but we did not find the same for *P. agricolaris* density (Figure 3). This difference between *P. agricolaris* and *P. hayleyella* is consistent with prior studies that found that *P. hayleyella* is more harmful than *P. agricolaris* (Shu *et al*., 2018a; Haselkorn *et al*., 2019; Khojandi *et al*., 2019). In addition to being more harmful, *P. hayleyella* also has a reduced genome relative to *P. agricolaris* (Brock *et al*., 2020; Noh *et al*., 2022). Reduced genomes are a common result of persistent host association in beneficial symbionts and pathogens (McCutcheon & Moran, 2012). While *P. hayleyella* has maintained its ability to harm hosts over this persistent association, it is important to remember that infection also comes with the ability to carry prey bacteria (DiSalvo *et al*., 2015; Khojandi *et al*., 2019). Thus, this symbiosis involves a balance between harm and benefits to hosts.

Whether a specific symbiosis involves fitness alignment or conflict may depend on a third party that affects the costs and benefits of symbiosis. Our results show that third parties can have complex effects on conflict and fitness alignment in symbioses; the symbiosis between *D. discoideum* and *Paraburkholderia* appears to involve elements of conflict and cooperation that depends on how third-party bacteria are eaten, carried, and left behind.

## Acknowledgements

This material is based upon work supported by the National Science Foundation under grants DEB-1753743, IOS-1656756, and DEB-2237266. We thank Tyler Larsen for comments on an early draft of the manuscript. We also thank Debbie Brock and the rest of the Strassmann/Queller lab group for feedback during the development and execution of this project.

