## Supplemental Figures 1 & 2 for "Complex third-party effects in the *Dictyostelium*-*Paraburkholderia* symbiosis: prey bacteria that are eaten, carried, or left behind"

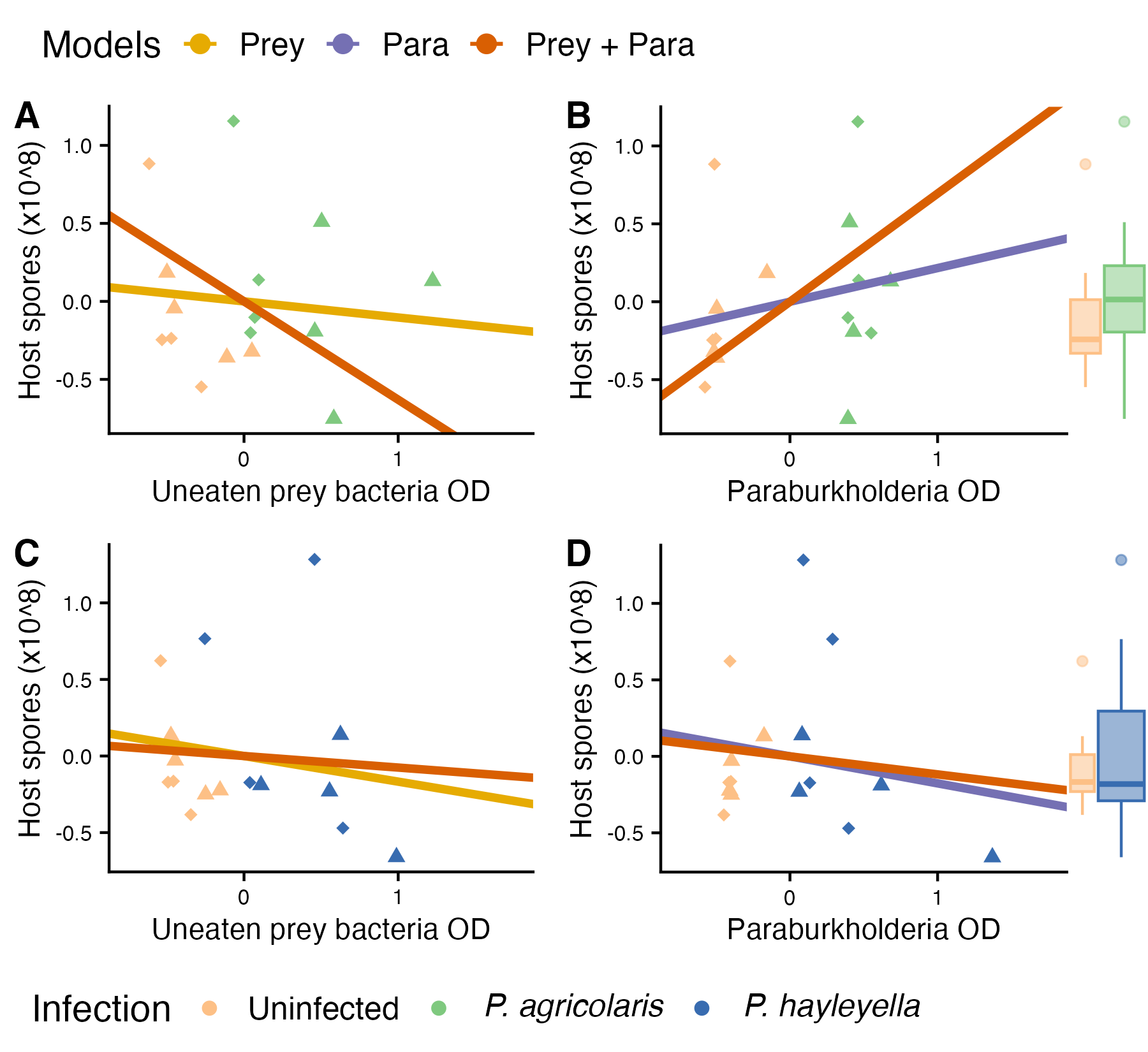

**Figure S1:** Host spore production for cured-and-reinfected hosts and uninfected controls as a function of uneaten food bacteria (A&C), *Paraburkholderia* density (B&D), and infection status (boxplots). Fit of models of host spore production from Figure 3A are shown as lines (models of continuous variables) or as boxplots (models for categorical infection status). All measurements were scaled as described in the methods.

Point shapes indicate different dates on which experiments were replicated.

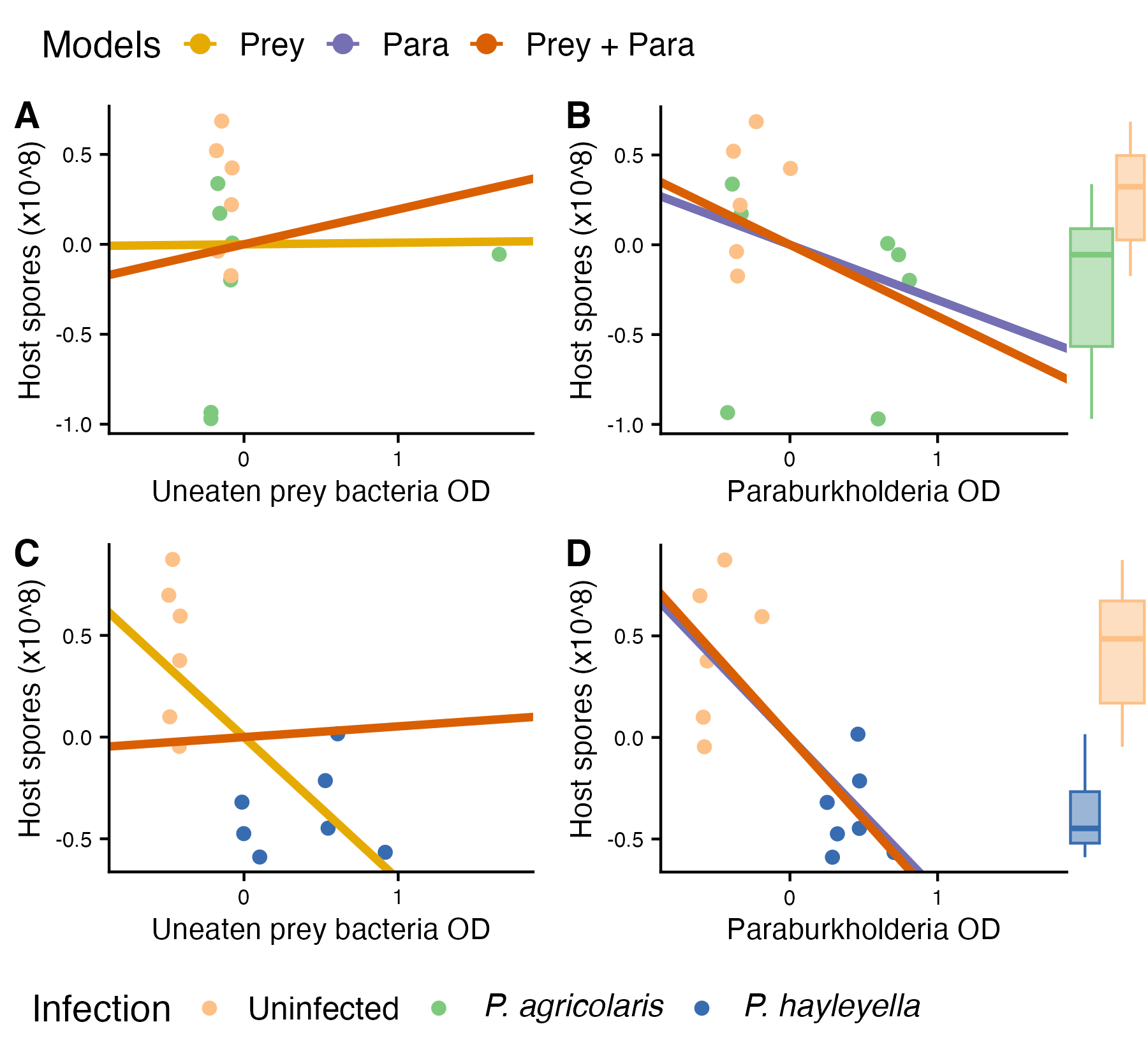

**Figure S2:** Host spore production for naturally infected hosts and uninfected controls as a function of uneaten food bacteria (A&C), *Paraburkholderia* density (B&D), and infection status (boxplots). Fit of models of host spore production from Figure 3B are shown as lines (models of continuous variables) or as boxplots (models for categorical infection status). All measurements were scaled as described in the methods.
